# In dystrophic *mdx* hindlimb muscles where fibrosis is limited versican haploinsufficiency transiently improves contractile function without decreasing inflammation

**DOI:** 10.1101/2024.05.07.592907

**Authors:** Danielle Debruin, Natasha L. McRae, Alex B. Addinsall, Daniel R. McCulloch, Robert G. Barker, Alan Hayes, Robyn M. Murphy, Nicole Stupka

## Abstract

The provisional matrix protein versican is upregulated in Duchenne muscular dystrophy. Versican heightens inflammation in fibrotic diseases and is involved in myogenesis. In fibrotic diaphragm muscles from dystrophic *mdx* mice, versican reduction attenuated macrophage infiltration and improved contractile function. We investigated the association between versican and *mdx* hindlimb muscle pathology, where inflammation and regeneration are increased but fibrosis is minimal. Immunohistochemistry and qRT-PCR were used to assess how fiber type and glucocorticoids (α-methylprednisolone) modulate versican expression. Female *mdx* and male versican haploinsufficient (hdf) mice were bred resulting in male *mdx*-hdf and *mdx* (control) pups. Versican expression, contractile function, and pathology were evaluated in fast extensor digitorum longus (EDL) and slow soleus muscles, excised under medetomidine-midazolam-fentanyl anesthesia.

Versican immunoreactivity was highest in soleus muscles. *Versican* mRNA transcripts were reduced by α-methylprednisolone in soleus, but not EDL, muscles. Versican expression was decreased in soleus muscles from 6-week-old *mdx*-hdf mice leading to increased force output and a modest reduction in fatiguability. These functional benefits were not accompanied by decreased inflammation; muscle architecture, regeneration markers, and fiber type also did not differ between genotypes. Improvements in soleus function were lost in adult (20-week-old) *mdx*-hdf mice with no significant effect of versican haploinsufficiency on macrophage infiltration and regeneration markers.

Soleus muscles from juvenile *mdx* mice were most responsive to pharmacological or genetic approaches targeting versican; however, the benefits of versican reduction were limited due to low fibrosis. Pre-clinical matrix research in dystrophy should account for muscle phenotype and the interdependence between the fibrosis and inflammation.

**NEW & NOTEWORTHY:** The proteoglycan versican is upregulated in muscular dystrophy. In fibrotic diaphragm muscles from *mdx* mice, versican reduction attenuated macrophage infiltration and improved performance. Here, in hindlimb muscles from 6- and 20-week-old *mdx* mice, where pathology is mild, versican reduction did not decrease inflammation and contractile function improvements were limited to juvenile mice. In dystrophic *mdx* muscles, the association between versican and inflammation is mediated by fibrosis, demonstrating interdependence between the immune system and extracellular matrix.

## INTRODUCTION

Duchenne Muscular Dystrophy (DMD) is an X-linked, pediatric disease caused by mutations in the dystrophin (*DMD*) gene. The loss of a functional dystrophin protein renders muscles highly susceptible to chronic damage and heightened inflammation, ultimately leading to the replacement of contractile tissue with fatty, fibrotic infiltrates (1). In patients with DMD, muscle regenerative capacity is also reduced (2). The interdependence between the immune system and the extracellular matrix is increasingly recognized as significant in disease pathology and tissue repair (3). In dystrophic muscles, fibrosis via feedforward mechanisms heightens inflammation, driving further extracellular matrix (ECM) expansion, precipitating muscle degeneration, as well as creating a hostile microenvironment for muscle regeneration (3, 4). A greater understanding of how dysregulated ECM components in dystrophic muscles contribute to disease progression is important for novel insight into disease pathology and the discovery of candidate therapeutics for DMD.

Fibrosis in DMD is characterized by increased muscle collagen content, as well as the persistent upregulation of various provisional matrix proteins and proteoglycans including versican (5–8). Versican is also upregulated in *mdx* mice, the most widely used mouse model of DMD (6, 9, 10). Excess versican is linked to fibrosis and inflammation in various organ systems, including the heart, lungs, and liver (11–14). Versican amplifies inflammatory responses by interacting with immune cell membrane proteins (e.g., CD44 and integrin-β1) to guide leukocyte trafficking (15). Versican also binds cytokines and chemokines, such as monocyte chemoattractant protein-1 (MCP-1) and TGFβ, via its chondroitin sulfate side chains to establish gradients which modulate cell behavior and signaling (15–18). Conversely, versican synthesis is stimulated by TGFẞ (19, 20) and in dystrophic muscles, TGFβ is an important pro-fibrotic molecule implicated in failed regeneration (1, 21). Versican is proteolytically processed by A Distintegrin and Metalloproteinase with Thrombospondin Motifs (ADAMTS) versicanases into versikine — a pro-inflammatory matrikine (22, 23). In dystrophic muscles, versican and versikine are co-localised to regions of muscle degeneration and inflammation, as well as muscle regeneration (6, 24). The provisional matrix regulates cellular processes relevant to skeletal muscle development and regeneration (25). The transient synthesis and subsequent degradation of versican by ADAMTS- 5 has an emerging role in myogenesis (26–28). *In vitro*, excess versican impairs the fusion of C2C12 myoblasts into multinucleated myotubes (7, 26).

Whilst various ECM components are dysregulated in dystrophic muscles, we propose that versican is highly relevant to DMD muscle pathology due to its role in inflammation, fibrosis and myogenesis, as well as its close association with TGFẞ. Glucocorticoids are the pharmacological standard of care for DMD (29), and have been shown to downregulate TGFβ signaling networks in hindlimb muscles from *mdx* mice (21). Glucocorticoids also decrease versican expression in various tissues and cells, including fetal lungs (30), lung fibroblasts (31), and C2C12 myoblasts (32). Whether glucocorticoids have a similar effect on dystrophic muscles *in vivo* is unknown.

We have previously used a genetic approach to test the hypothesis that versican reduction would ameliorate dystrophic muscle pathology (24). Specifically, female C57BL/10 *mdx* mice were bred with male heart defect (hdf) mice which are haploinsufficient for the versican allele (33). In diaphragm muscles from the adult *mdx*-hdf offspring, versican reduction decreased macrophage infiltration and improved *ex vivo* strength and endurance compared with diaphragm muscles from *mdx* littermate controls (24). In *mdx* mice, the diaphragm best models the fibrotic aspect of DMD pathology, including an upregulation of versican (32, 34). These findings showed that versican is a pro-inflammatory ECM component in dystrophic muscles. However, DMD is a pediatric disease occurring on a background of growth, which involves myonuclear accretion and myofiber hypertrophy (35) and encompasses extensive ECM remodeling (36). Therefore, we investigated the effects of versican haploinsufficiency on the pathology and contractile function of slow soleus and fast extensor digitorum longus (EDL) hindlimb muscles from juvenile and adult (e.g., 6- and 20-week-old) *mdx*-hdf mice and *mdx* control littermates. In hindlimb muscles from *mdx* (C57BL/10ScSn^mdx^) mice, necrosis peaks at 3-4 weeks of age and is followed by effective regeneration and stabilization of pathology by 8 weeks of age (37). Puberty occurs between 5-7 weeks of age and corresponds to a decrease in growth rate; however, muscle damage, myonecrosis and regeneration persist into adulthood – albeit with minimal fibrosis – with approximately 60% of adult *mdx* myofibers affected at any one time (38). Hence, the 20- week time point would allow for the assessment of the longer-term consequences of versican haploinsufficiency on muscle pathology.

## MATERIALS AND METHODS

### Animal studies

To assess the effect of versican haploinsufficiency on the pathology of *mdx* hindlimb muscles, female *mdx* (C57BL/10ScSn^mdx^) mice obtained from the Animal Resource Centre (Australia), were bred with male hdf mice, haploinsufficient for the versican allele (hdf) obtained from Hoffman-La Roche (USA) (39). The resulting F1 *mdx* and *mdx*-hdf male pups were confirmed through genotyping and demonstrated the expected Mendelian genetic ratios (24). Juvenile and adult mice were maintained in grouped cages (2–5 mice per cage) until 6- or 20-weeks of age. These experiments were approved by the Deakin University Animal Ethics Committee (projects A79/2011 and G06/2015)

Hindlimb muscle samples from *mdx* (C57BL/10ScSn^mdx^) were used to investigate how versican expression is affected by muscle fiber type composition, age, and glucocorticoid (α- methylprednisolone) treatment. EDL and soleus muscles were obtained from juvenile *mdx* mice that received α-methylprednisolone from day 18 to day 28 postnatum, as previously described (40). In addition, soleus, tibialis anterior and EDL muscle samples were sourced from 6- week and 20-week-old untreated *mdx* mice. This study was approved by the Victoria University Animal Experimental Ethics Committee (project 19003).

### Ex vivo contractile function testing of hindlimb muscles from mdx and mdx-hdf mice

The contractile function experiments were conducted as previously described (41–43). Briefly, mice were deeply anaesthetized via an intraperitoneal injection of medetomidine (0.6 mg/kg), midazolam (5 mg/kg) and fentanyl (0.05 mg/kg). The proximal and distal tendons of EDL or soleus muscles were exposed and tied with 5.0 suture silk. The muscles were then excised tendon to tendon and placed in an organ bath containing Kreb’s ringer buffer (NaCl 137 mM, NaHCO_3_ 24 mM, D-glucose 11 mM, KCl 5 mM, NaH_2_PO_4_·H_2_O 1mM, MgSO_4_ 1 mM, CaCl_2_ 2 mM, d-tubocurarine chloride 0.025 mM) maintained at 25 °C and bubbled with Carbogen (95 % CO_2_ and 5 % O_2_). The distal tendon of the muscle was tied to an immobile pin and the proximal tendon was attached to the lever arm of a dual mode force transducer (300-CLR; Aurora Scientific). Following determination of optimum muscle length (L_o_), the maximal force producing capacity was determined via a force frequency curve, ranging from 1 to 120 Hz (350 ms train duration for EDL and 800 ms for soleus). To evaluate fatigability, repeated, intermittent stimuli (60 Hz) were administered every 5 seconds for 4 minutes. Force recovery was monitored at 2-, 5- and 10- minutes post fatigue (60 Hz). (41–43). All stimulation parameters and contractile responses were controlled and recorded using Dynamic Muscle Control Software (DMCv5.415) along with force transducer control/feedback hardware (Aurora Scientific). Force output was analyzed using Dynamic Muscle Analysis Software (Aurora Scientific). Muscle mass was recorded following the contractile function protocol. Overall muscle cross-sectional area was determined by dividing muscle mass by the product of optimum fiber length (L_f_) and 1.06 mg·mm^3^ (density of mammalian muscle). L_f_ for the soleus is 0.71 x L_o_ and L_f_ for the EDL is 0.44 x L_o_. Force output (P_o_) was normalized to muscle cross-sectional area to obtain specific force output (sP_o_). The force frequency curve data were also normalized to maximal force output to assess a potential fiber type shift (44). The fatigue and recovery data were presented as the percentage of the initial force output.

### Tissue collection

EDL, soleus, and tibialis anterior muscles were snap frozen, with the contralateral hindlimb muscles mounted in optimal cutting temperature (OCT) medium and frozen in liquid-nitrogen cooled isopentane for histology and immunohistochemistry.

### Histology for muscle pathology, fibrosis, and oxidative capacity

Hindlimb muscle cryosections (8 µm thick) were stained with hematoxylin and eosin (H&E) to assess general muscle morphology or with picrosirius red (PSR) to stain for collagen. For the H&E stained soleus and EDL muscle cross-sections from *mdx*-hdf and *mdx* mice, Image-Pro Plus software (version 7; Media Cybernetics) was used to measure the myofiber size (expressed as min ferret diameter), centrally nucleated myofibers, and the percentage of muscle cross-section comprised of basophilic, mononuclear infiltrates and degeneration (45, 46).

Frozen TA muscle were cryosectioned and stained within 24 h for succinate dehydrogenase (SDH) activity to assess oxidative capacity as previously described (47). Following staining, samples were imaged on a slide scanner microscope (ZEISS Axio Imager Z2) and captured using x20 objective. For analysis, the image of the whole TA cross-section was converted to an 8-bit grayscale format and the staining intensity of both the oxidative and glycolytic regions of the TA was analyzed by thresholding using ImageJ software (NIH).

### Immunohistochemistry for versican, versikine, CD68+ macrophages and desmin+ muscle progenitor cells

Immunohistochemistry for V0/V1 versican (anti-GAGβ; Millipore, AB1033), versikine (anti- DPEAAE neo-epitope; ThermoFisher Scientific, PA1-1748A) (24, 26, 48, 49), anti-CD68 (Abcam, Ab125212), and desmin (Abcam, Ab15200) (24) was performed as previously described. For the V0/V1 versican and versikine stained muscle cross-sections, an average of three non- overlapping representative digital images were captured with a confocal microscope of each muscle cross-section (Olympus Fluoview FV10i). To determine the percentage of muscle cross- section immunoreactive for versican or versikine, planimetric analysis of the digital images was completed using Image-Pro Plus software. For analysis of CD68 positive monocytes and macrophages or desmin positive muscle progenitor cells, digital images of the whole muscle cross-section were captured with an IX81 Olympus fluorescence microscope and ImageJ software was used to determine the number of positive cells per mm^2^ of tissue cross-section.

### Immunohistochemistry for muscle fiber type

For muscle fiber typing soleus and TA cross-sections were reacted with a primary antibody cocktail containing, anti-myosin heavy chain IIa (MyHC IIa) (Developmental Studies Hybridoma Bank (DSHB); SC-71), anti-MyHC IIb (DSHB; BF-F3), and anti-MyHC I (DSHB; BA-F8) as described by Bloemberg *et al.* (50). Digital images of the whole muscle cross-section were then captured with an IX81 Olympus fluorescence microscope and ImageJ software was used to count the number of type I, IIa and IIb positive fibers per mm^2^ of tissue cross-section.

### RT-qPCR for gene expression analysis

EDL, soleus, and TA muscles were homogenized in 1 mL of TRIzol agent (Thermo Fisher Scientific) prior to RNA extraction (RNeasy kit; Qiagen). Thereafter, methods of reverse transcription (iScript cDNA synthesis kit; Biorad), Real Time quantitative PCR (IQ SYBR Green super mix; Bio-Rad) and cDNA quantitation (Quant-iT OliGreen ssDNA reagent kit; Invitrogen) were followed as previously described (32). Relative changes in mRNA levels were calculated using the ΔCt method, with real time data normalized to cDNA content. The average of the control group was set to 1. Oligonucleotide primer sequences for the genes of interest are shown in Table 1.

**Table 1.**
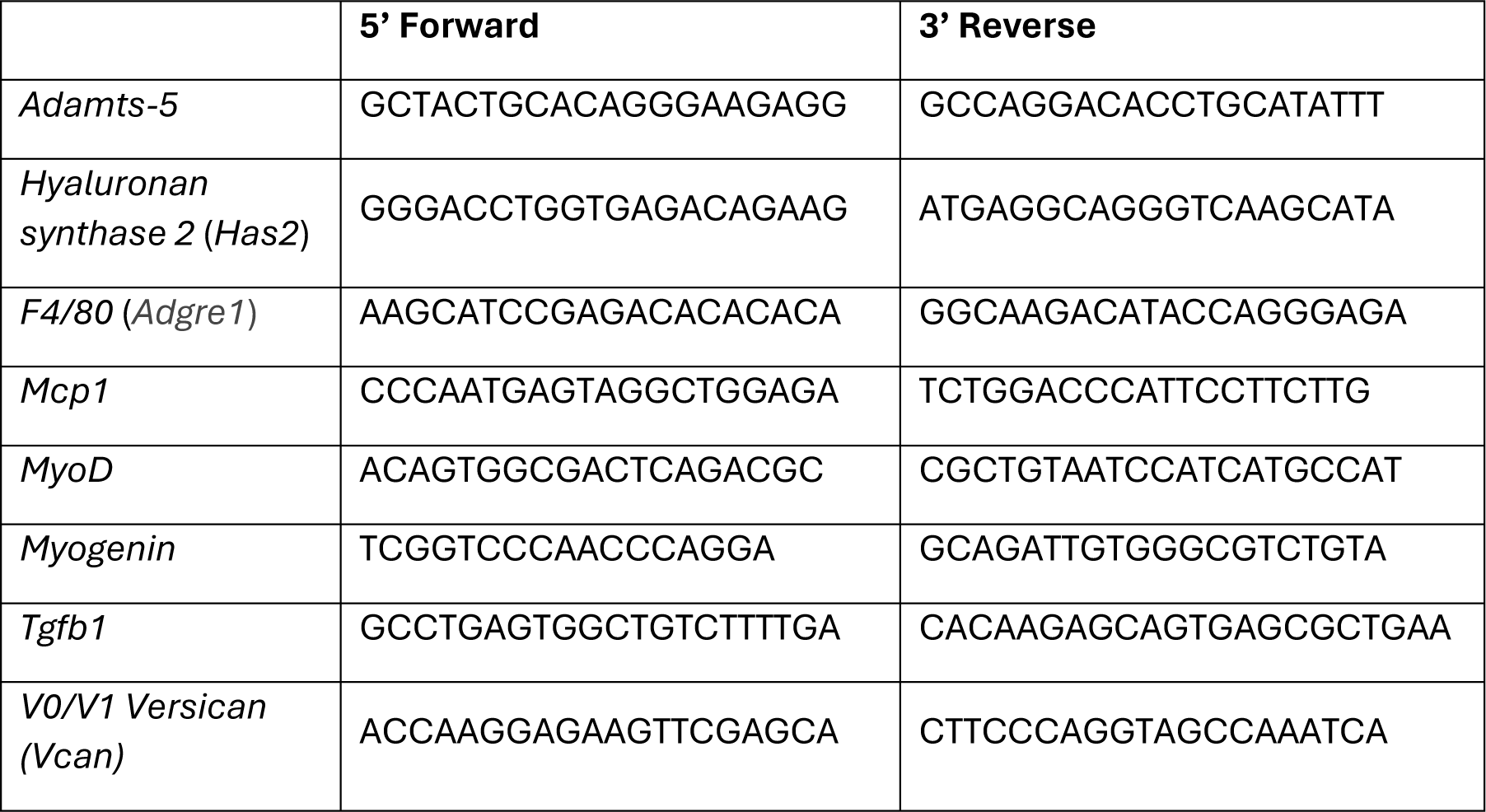
Primer sets used for RT-qPCR.

### Statistics

An independent sample T-test was undertaken for all histology, immunohistochemistry, and gene expression analyses. To assess muscle strength and fatigability, a 2-way ANOVA with the factors being genotype and stimulation frequency (Hz) or time was performed and followed by Tukey’s post hoc analysis where appropriate. All data are presented as mean ± S.E.M. and the alpha value was set to 0.05.

## RESULTS

### Phenotype dependent differences in versican expression and degradation in mdx hindlimb muscles

H&E-stained soleus, tibialis anterior and EDL muscle cross-sections from 6- and 20-week-old *mdx* mice presented with inflammatory, basophilic foci and centrally nuclei fibers indicative of recent damage and repair and with a very low level of fibrosis as determined by collagen staining (Fig. 1). Despite limited fibrosis, versican, versikine and ADAMTS-5 were localized to the endomysium of all hindlimb muscles. The presence of versikine demonstrated ongoing versican degradation by ADAMTS versicanases such as ADAMTS-5 (51), which is highly expressed in dystrophic muscles (41, 52). Slow soleus muscles from 6-week-old *mdx* mice appeared to have greater versican, versikine and ADAMTS-5 immunoreactivity versus fast TA and EDL muscles across both age groups.

**Figure 1.**
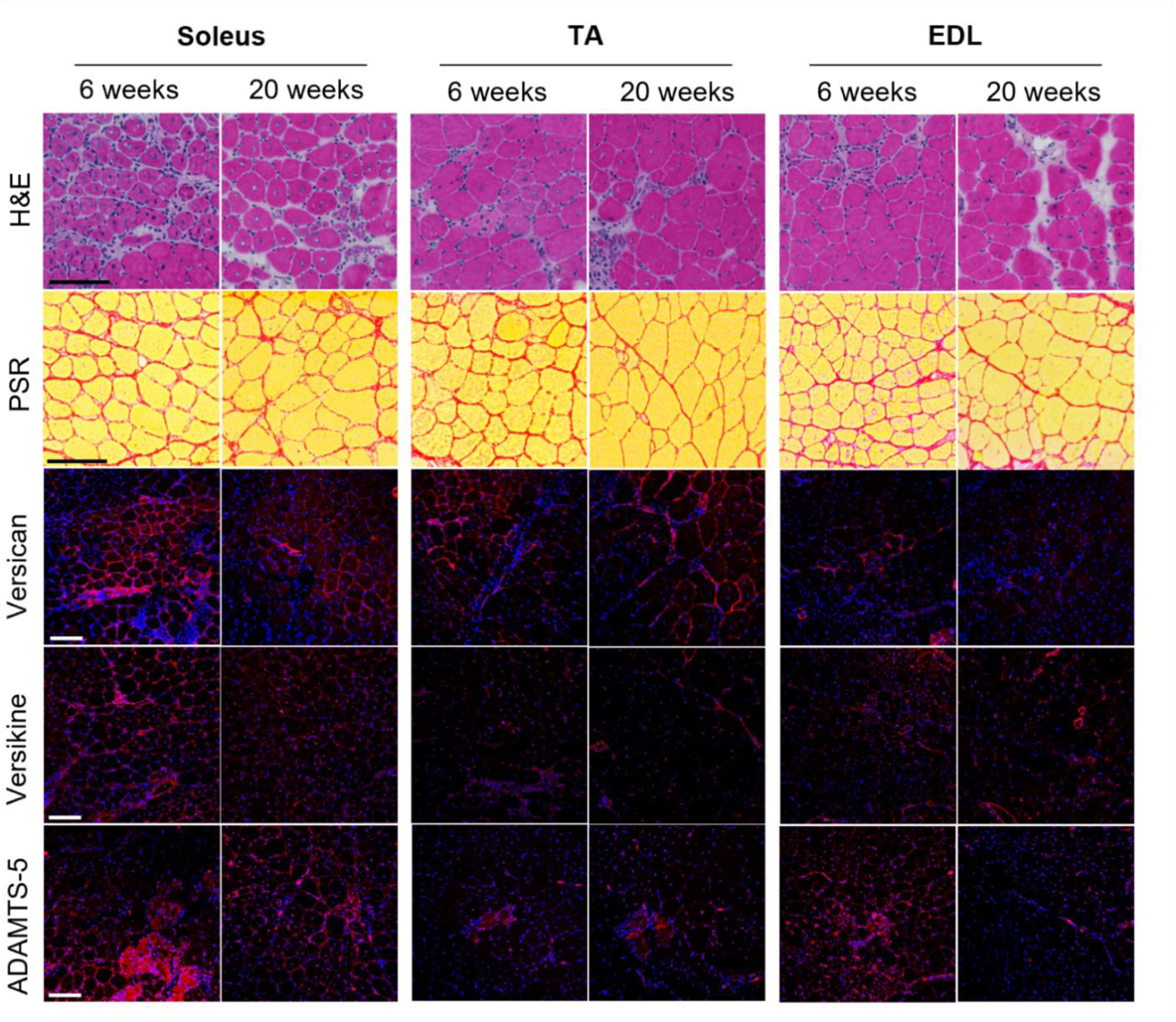
Muscle pathology in slow (soleus) and fast tibialis anterior (TA) and extensor digitorum longus (EDL) hindlimb muscles from *mdx* mice at 6- and 20-weeks of age. H&E for muscle architecture. Picrosirius red (PSR) for fibrosis with collagen staining red and muscle fibers yellow. Versican, versikine or ADAMTS-5 immunoreactivity (red) with a DAPI nuclear counterstain (blue). Scale bar = 100 μm; *N* = 3-4

### Tgfb1 and V0/V1 Versican mRNA transcript abundance in mdx hindlimb muscles following α- methylprednisolone treatment

The effect of α-methylprednisolone on *Tgfb1* and *V0/V1 Versican* gene expression was investigated in hindlimb muscles from juvenile *mdx* mice, given that these target genes are closely associated and downregulated by glucocorticoids in C2C12 myotubes (9, 18–20). The association between versican and TGFẞ was more evident in soleus than EDL muscles. Specifically, *Tgfb1* (*P* < 0.01) and *V0/V1 Versican* (*P* < 0.01) mRNA transcripts were decreased in soleus muscles following 10 days of treatment with α-methylprednisolone beginning at 18 days of age, a period that corresponds with peak hindlimb muscle necrosis (40). Whereas, in EDL muscles, α-methylprednisolone increased *Tgfb1* gene expression (*P* < 0.01), with no significant effect on *V0/V1 Versican* mRNA transcripts (Fig. 2A and B).

**Figure 2.**
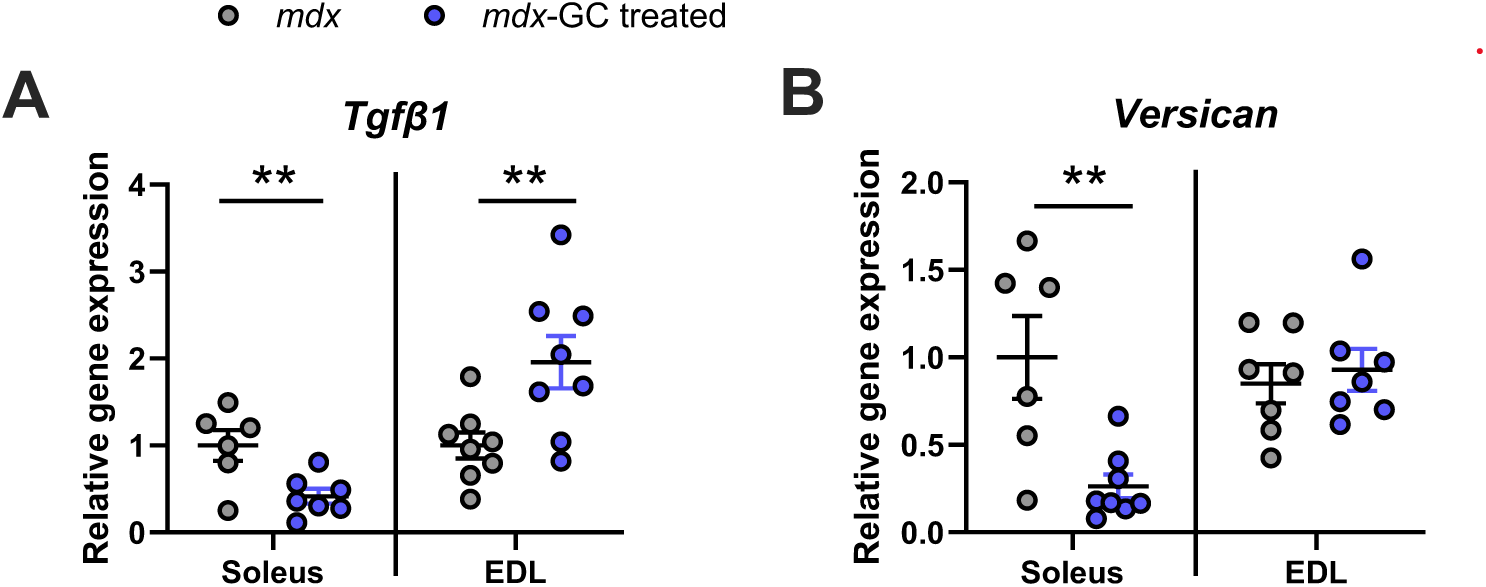
Effect of glucocorticoids (GC) on *Tgfβ1* and *V0/V1 Versican* gene expression in dystrophic hindlimb muscles. With α-methylprednisolone treatment, (A) Tgfβ1 mRNA transcripts were increased in EDL and decreased in soleus (SOL) muscles; whilst (B) *V0/V1 Versican* mRNA transcripts were decreased in soleus muscles only. **P < 0.01; independent T-test. *N* = 6-8.

These initial characterization experiments indicate that in *mdx* hindlimb muscles the expression of versican and its remodeling into the bioactive versikine fragment is phenotype-dependent and appears to be more dynamic in slow soleus muscles.

### Versican haploinsufficiency and versican expression in hindlimb muscles from 6-week-old mdx mice

A trend for reduced *V0/V1 versican* mRNA transcript abundance was observed in soleus muscles from *mdx*-hdf mice (Fig. 3A; *P* = 0.09). *V0/V1 versican* gene expression was reduced by approximately 50% in TA muscles from *mdx*-hdf mice compared to *mdx* littermates (Fig. 3B; P < 0.05), but not EDL muscles (Fig. 3C; *P* = 0.248). Irrespective of muscle phenotype, the genetic reduction of versican had no significant effect on the mRNA transcript abundance of *Adamts-5* nor *Hyaluronan synthase-2* (*Has-2*), the predominant Has isoform responsible for the synthesis of hyaluronan, provisional matrix component and a key versican binding partner in skeletal muscle (11, 53). This suggests that the transcriptional regulation of these key provisional matrix genes is independent of versican haploinsufficiency (Fig. 3D-I).

**Figure 3.**
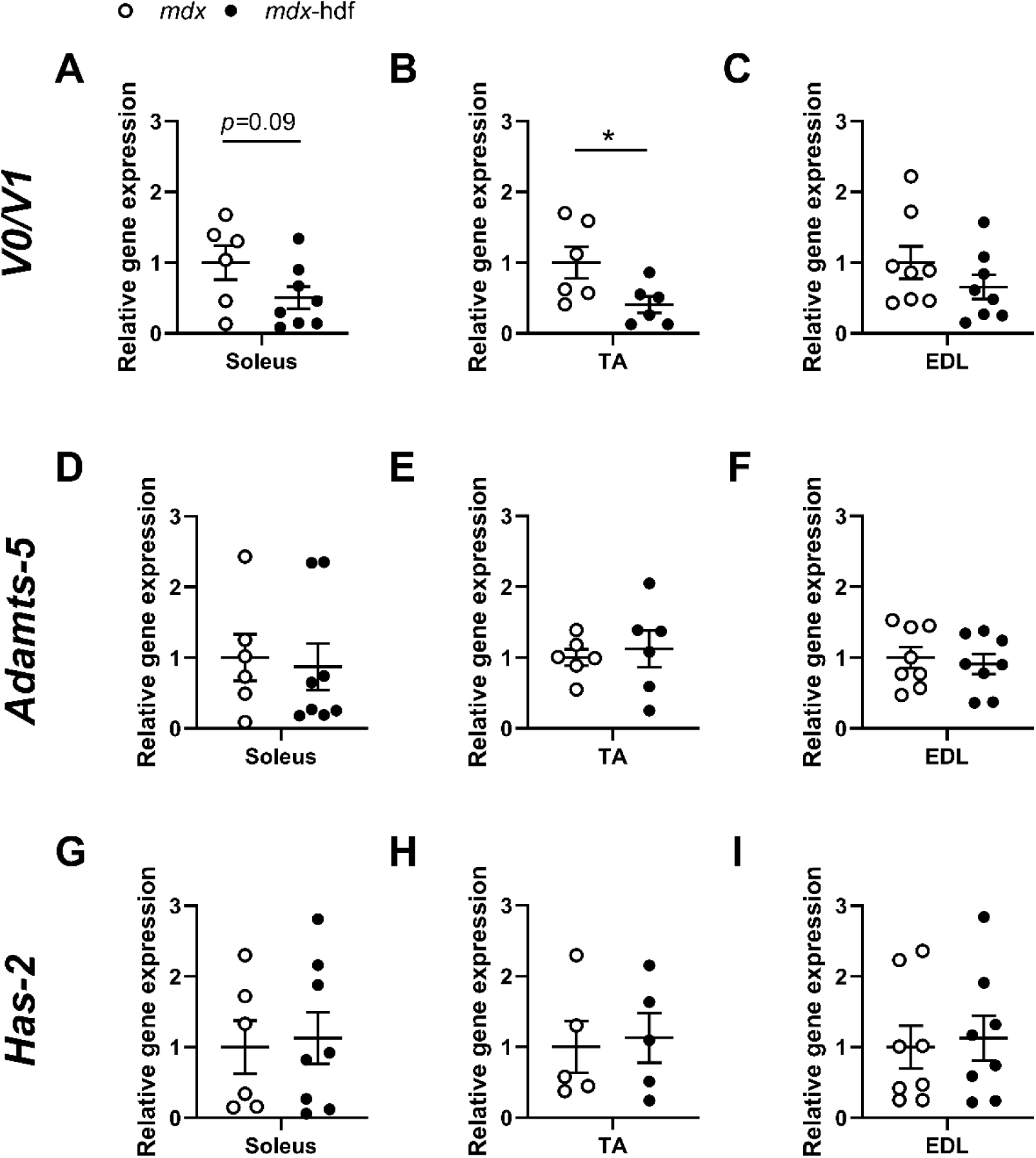
Provisional matrix gene expression between the extensor digitorum longus (EDL), soleus (SOL) and tibialis anterior (TA) muscles of 6-week-old *mdx* and *mdx-*hdf mice. (A-C) *V0/V1 Versican*, (D-E) *Adamts-5* and (G-I) *Hyaluronan synthase-2* (*Has-2*) gene expression in hindlimb muscles from *mdx*-hdf mice and littermate controls. *P < 0.05; independent T-test. *N =* 6-8 per genotype.

The protein expression of full length V0/V1 versican and versikine was determined in soleus and EDL muscles from *mdx*-hdf and *mdx* mice using immunohistochemistry (Fig. 4), as these hindlimb muscles were used for the assessment of *ex vivo* contractile function. V0/V1 versican immunoreactivity was reduced in soleus muscles from *mdx*-hdf mice when compared to *mdx* littermate controls (P < 0.01; Fig. 4A); however, no significant difference in versikine immunoreactivity was observed (Fig. 4B). The effect of versican haploinsufficiency on decreasing V0/V1 versican, but not versikine, immunoreactivity was previously observed in diaphragm muscles from *mdx-*hdf mice (24). Versikine can be further degraded by ADAMTS versicanases, released into circulation, or retained in the matrix (51). The regulation of versikine in dystrophic muscles is currently not well understood.

**Figure 4.**
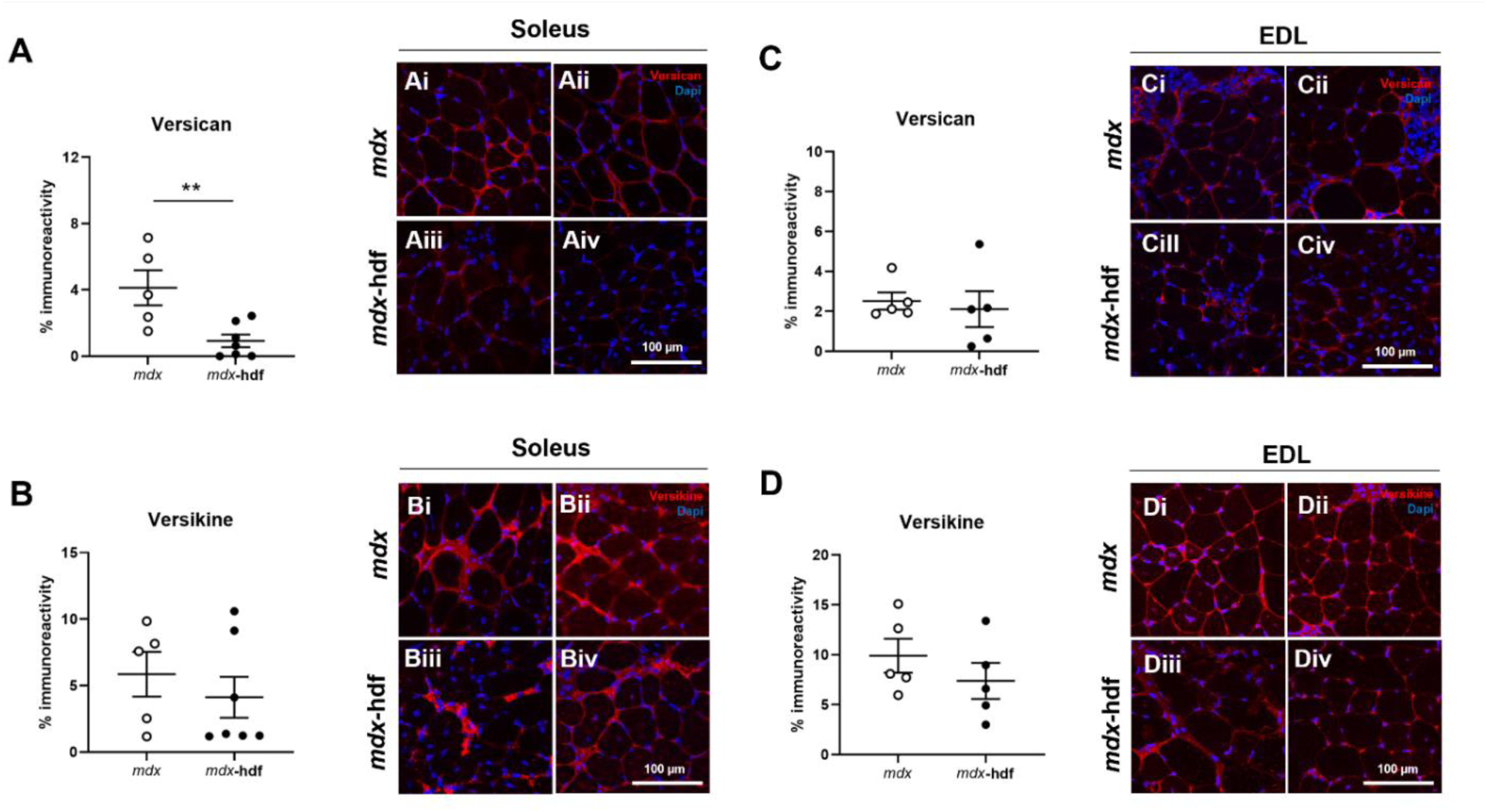
V0/V1 versican and versikine immunoreactivity in soleus and EDL muscles from 6-week-old *mdx* and *mdx*-hdf mice. Quantification of versican and versikine immunoreactivity in soleus (A and B) and EDL (C and D) muscles. Representative images of versican (Ai-iv and Ci-iv) and versikine (Bi-iv and Di-iv) staining of soleus and EDL muscles, respectively, from three different *mdx* and *mdx*-hdf mice. Versican or versikine immunoreactivity (red) with a DAPI nuclear counterstain (blue). Scale bar = 100 μm. *N =* 5-7 per genotype. **P < 0.01; independent T-test. *N =* 6-8 per group.

Whereas, in EDL muscles from *mdx*-hdf and *mdx* mice, V0/V1 versican (Fig. 4C) or versikine (Fig. 4D) immunoreactivity was not significantly different. This lack of effect of versican haploinsufficiency on decreasing V0/V1 versican immunoreactivity in EDL versus soleus or diaphragm muscles (24), may be due to muscle specific differences in fiber type dependent ECM content or dystrophic pathology (34, 54).

### Versican haploinsufficiency has fiber type specific effects on the ex vivo contractile properties of hindlimb muscles from 6-week-old mdx mice

Body weight and the absolute or normalized to body weight mass of soleus and EDL muscles did not significantly differ between *mdx*-hdf mice and *mdx* littermates (Fig. 5A-C). *Ex vivo* force output was determined by conducting a force frequency curve. Soleus muscles from *mdx*-hdf mice produced higher absolute forces across increasing frequencies compared to *mdx* littermates (P < 0.0001; Fig. 5D). When force output was normalized to muscle cross-sectional area, an upward shift in the specific (sP_o_) force frequency curve was observed in soleus muscle from *mdx*-hdf mice (P < 0.001; Fig. 5E). Thus, versican reduction has favorable effects on soleus muscle force producing capacity in juvenile *mdx* mice. When soleus force frequency curve output values were expressed as a percentage of maximal force, no left or right axis shifts were observed (Fig. 5F). This suggests a lack of effect of versican haploinsufficiency on intracellular calcium dynamics or muscle fiber type composition (44). There was no significant difference in the absolute (P_o_) or specific force (sP_o_) force output of the EDL muscles from *mdx*-hdf mice and *mdx* littermate controls across increasing stimulation frequencies (Fig. 5G and H), nor was a left or right axis shift was observed (Fig. 5I). Following determination of maximal force output, the effect of versican haploinsufficiency on muscle fatiguability was assessed. Following 4 minutes of intermittent stimulation, a modest reduction in fatigability was observed in soleus muscles from *mdx*-hdf mice compared to *mdx* littermates (P < 0.001; Fig. 5J); although, force recovery did not significantly differ between genotypes. EDL muscles from *mdx*-hdf mice fatigued less than EDL muscles from *mdx* littermates (P < 0.05, Fig. K). Force recovery was also improved with versican haploinsufficiency (P < 0.05; Fig. 5K). Therefore, in dystrophic hindlimb muscles, versican haploinsufficiency improved the *ex vivo* strength of soleus muscles and had greater positive effects on the fatiguability of EDL versus the soleus, despite no significant decrease in versican expression at 6 weeks of age.

**Figure 5.**
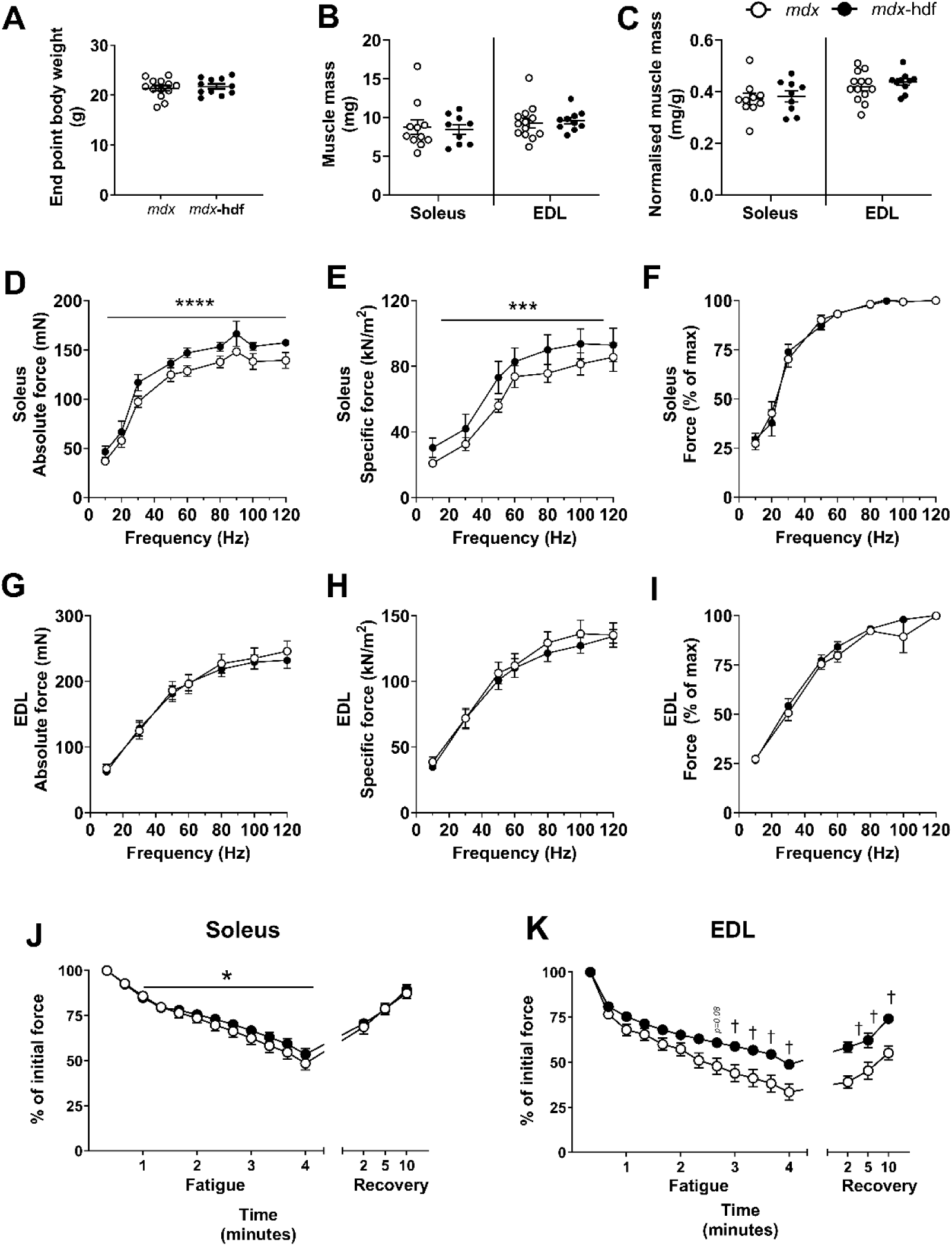
*Ex vivo* strength and endurance of soleus and EDL muscles from 6-week-old *mdx* and *mdx*-hdf mice. Endpoint body weight (A) and mass of soleus and EDL muscles expressed as (B) absolute mass and (C) normalized to body weight (mg of muscle mass per g of body weight). Force frequency curves of the soleus expressed as (D) absolute force (P_o_), (E) specific force normalized to muscle cross-sectional area (sP_o_), and as (F) a percentage of maximal force output. Force frequency curves of the EDL expressed as (G) absolute force, (H) specific force and as (I) a percentage of maximal force output. Soleus (J) and EDL (K) muscle fatigability and force recovery following 4 min of intermittent stimulation (60 Hz, every 5 s) expressed as a percentage of initial force output. *P < 0.05, *\*\*\**P *<* 0.001; *\*\*\*\**P *<* 0.0001; 2-way ANOVA – main effect genotype. ^+^P < 0.05; 2-way ANOVA – genotype x time interaction. *N =* 11-13 per group.

### Versican haploinsufficiency does not ameliorate the pathology of hindlimb muscles from 6- week-old mdx mice

The myofiber size distribution did not significantly differ between soleus or EDL muscles from *mdx*-hdf mice and *mdx* littermate controls, indicating that versican haploinsufficiency did not influence myofiber size (Fig. 6C and D). This is consistent with the lack of changes in muscle mass (Fig. 5B). Versican haploinsufficiency also had no significant effect on the percentage of muscle cross-section comprised of basophilic, mononuclear infiltrates and degenerating tissue (Fig. 6E). Versican synthesis and proteolysis is regulated during muscle development (26–28), and hence proportion of damaged and regenerated, centrally nucleated myofibers and the gene expression of the muscle-specific basic-helix-loop-helix transcription factors *Myod* and *Myogenin* were assessed. In soleus or EDL muscles from *mdx* and *mdx*-hdf mice, there was no significant difference in centrally nucleated myofibers (Fig. 6F) nor in the mRNA transcript abundance of *Myod* and *Myogenin* (Fig. 6G and H). Therefore, in juvenile *mdx* mice, versican haploinsufficiency does not appear to have a deleterious effect on postnatal growth or regenerative myogenesis.

**Figure 6.**
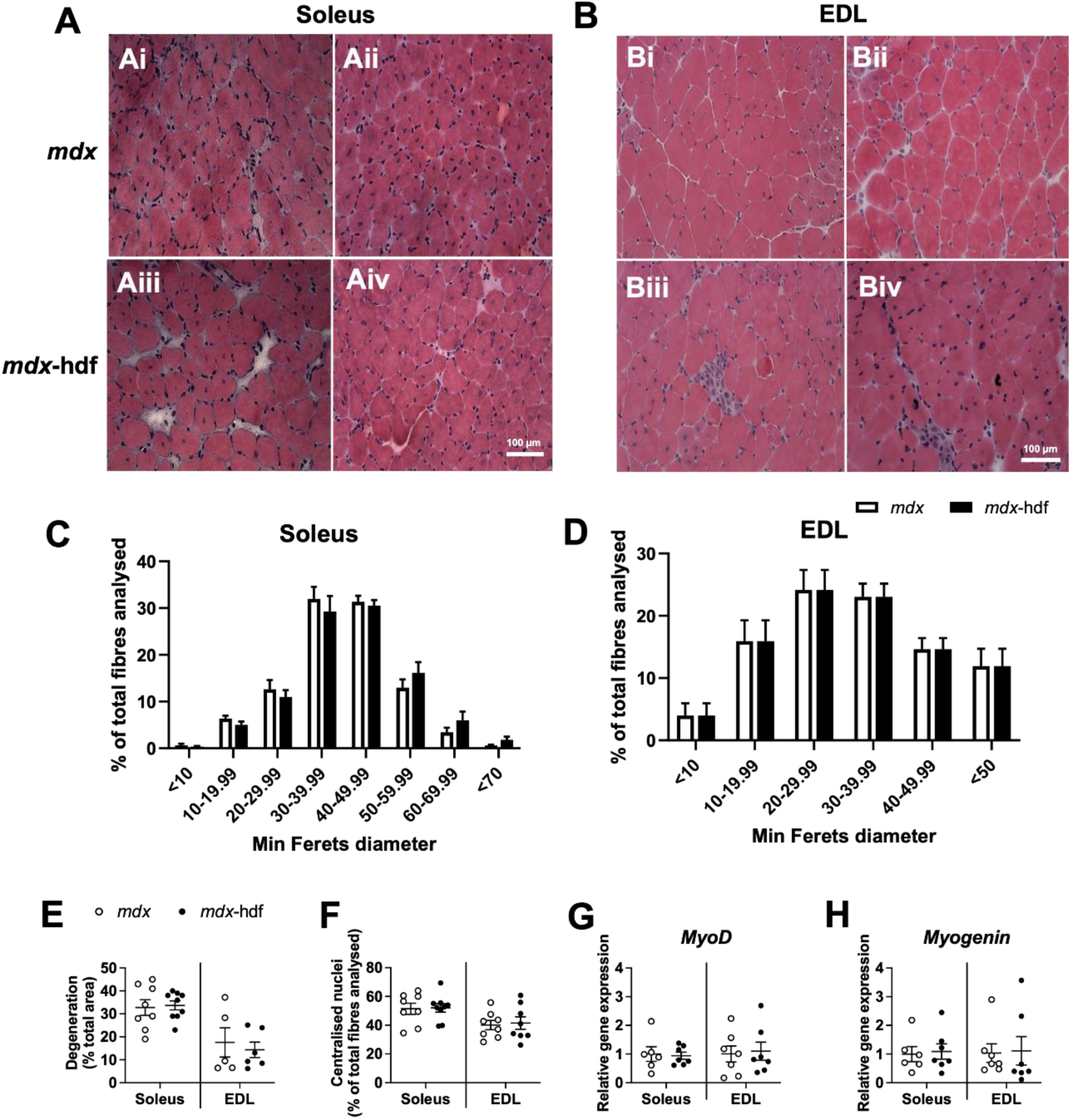
Evaluation of muscle morphology and gene markers of regenerative myogenesis in EDL and soleus muscles from 6-week-old *mdx-*hdf and *mdx* mice. Representative H&E-stained images for soleus (A) and EDL (B) muscles from *mdx* (Ai-ii; Bi-ii) and *mdx-*hdf (Aiii-iv; Biii-iv) mice. Myofiber size distribution in (C) soleus and (D) EDL muscles from *mdx-*hdf and *mdx* mice. (E) The percentage of muscle cross-section comprised of mononuclear infiltrates and degenerating tissue. (F) The percentage of centrally nucleated fibers in soleus and EDL muscles. (G, H) *MyoD* and *Myogenin* mRNA transcript abundance in soleus and EDL muscles. Scale bar = 100 μm. *N = 6* per group for histology and *N = 8* per group for gene expression analysis.

The number of CD68 positive monocytes and macrophage cells per mm^2^ of tissue cross-section did not significantly differ between soleus muscles from *mdx*-hdf mice and *mdx* littermate controls (Fig. 7A). The lack of effect on inflammation was corroborated by the observation that in both soleus and EDL muscles, the expression of pro-inflammatory and pro-fibrotic genes — *Tgfb1*, *Mcp1* and *F4/80* (also known as *Adgre1,* a pan-macrophage marker) — did not significantly differ between the *mdx*-hdf and *mdx* genotypes (Fig. 7B-D). Hence, the increase in force producing capacity of soleus muscles from 6-week-old *mdx*-hdf mice did not align to any of these observable changes in muscle pathology.

**Figure 7.**
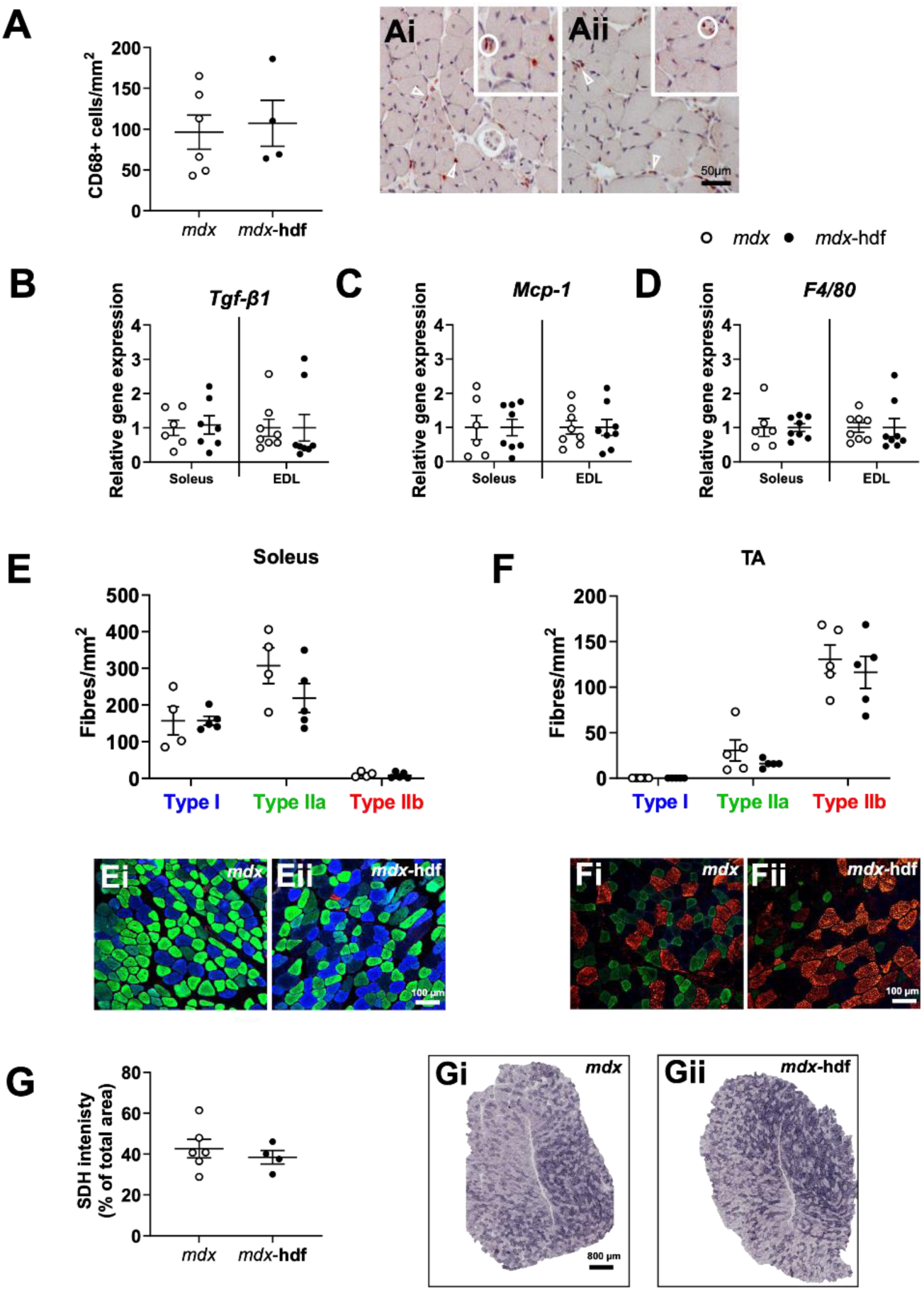
Effects of versican haploinsufficiency on inflammation and fiber type in hindlimb muscles from 6-week-old *mdx-*hdf and *mdx* mice. (A) The number of CD68 positive monocytes and macrophages per mm^2^ of tissue in soleus muscles from *mdx-*hdf mice and *mdx* littermate controls. Representative images of CD68 positive cells in soleus muscles from *mdx* (Ai) and *mdx-* hdf (Aii) mice. CD68 positive cells are stained reddish-brown with 3-amino-9-ethyl carbazole (AEC; white circle) and nuclei are counterstained with haematoxylin; scale bar = 50 μm. (B-D) *Tgfb1*, *Mcp1* and *Adgre1* (*F4/80*) mRNA transcript abundance in soleus and EDL muscles. The number of fibers expressing the myosin heavy chain (MyHC) type I, IIa or IIb isoforms per mm^2^ of muscle cross-section in (E) soleus and (F) tibialis anterior (TA) muscles. Representative images of MyHC staining in soleus and TA muscles from *mdx* (Ei & Fi) and *mdx-*hdf (Eii & Fii) mice with MyHC type I fibers stained blue, MyHC IIa fibers stained green, and MyHC IId fibers stained red; scale bar = 100 μm. (G) Succinate dehydrogenase (SDH) staining intensity in TA muscles. Representative images of SDH stained muscle cross-sections from *mdx* (Gi) and *mdx-*hdf mice (Gii) with darker staining indicative of fibers with a higher oxidative capacity. Scale bar = 800 μm. *N = 4-6* per group for immunohistochemistry or histology and *N = 8* per group for gene expression analysis.

As a reduction in muscle fatigability was observed in fast EDL muscles and to a lesser extent in slow soleus muscles from *mdx*-hdf mice (Fig. 5J and K), myosin heavy chain (MyHC) isoform immunostaining and succinate dehydrogenase (SDH) staining intensity – a histological marker of mitochondrial content – were to assess whether versican haploinsufficiency resulted in a shift to a slower muscle phenotype. Due to tissue limitations, TA, rather than EDL, muscles were used for these analyses as both muscles are of a fast phenotype and localized in the same myofascial compartment anatomically. MyHC I and IIa were the predominant isoforms expressed in soleus muscles and MyHC IIb was the predominant isoform expressed in TA muscles (50). The number of I, IIa or IIb MyHC positive fibers per mm^2^ muscle cross-section did not significantly differ between soleus or TA muscles from *mdx*-hdf mice and *mdx* littermate controls (Fig. 7E and F). Furthermore, in TA muscles, versican haploinsufficiency had no significant effect on SDH staining intensity (Fig. 7G). Therefore, a fiber type shift to a slower phenotype cannot account for the functional improvement in muscle fatiguability with versican haploinsufficiency.

### Versican haploinsufficiency exacerbates the pathology of soleus muscles from 20-week-old mdx mice

Hindlimb muscles from adult *mdx* mice experience ongoing cycles of damage and repair (38), with ECM synthesis and remodeling being integral to the inflammatory and regenerative response following injury (55, 56). Given that the genetic reduction of versican was more robust in soleus versus EDL muscles (Fig. 4), soleus muscles from 20-week-old *mdx*-hdf and *mdx* mice were used to assess the longer-term effects of versican haploinsufficiency on hindlimb muscle contractile function and pathology. The percentage of muscle cross-section immunoreactive for V0/V1 versican or versikine appeared to be lower in soleus muscles from adult 20-week-old compared to juvenile 6-week-old mdx mice (Fig. 4A and B; 8A and B). Indeed, in healthy skeletal muscle, versican is more highly expressed during development and postnatal growth versus adulthood (57). V0/V1 versican and versikine immunoreactivity did not significantly differ between soleus muscles from *mdx*-hdf mice versus *mdx* littermate controls (Fig. 8A and B). *V0/V1 versican* mRNA transcript abundance also did not differ between genotypes (P = 0.71; data not shown). The lack of effect of versican haploinsufficiency in soleus muscles from adult *mdx* mice is in contrast to our observations with soleus muscles from 6-week-old *mdx-*hdf mice (Fig. 4A and B) and our previously published findings in diaphragm muscles from adult *mdx-*hdf mice where fibrosis is high (24).

**Figure 8.**
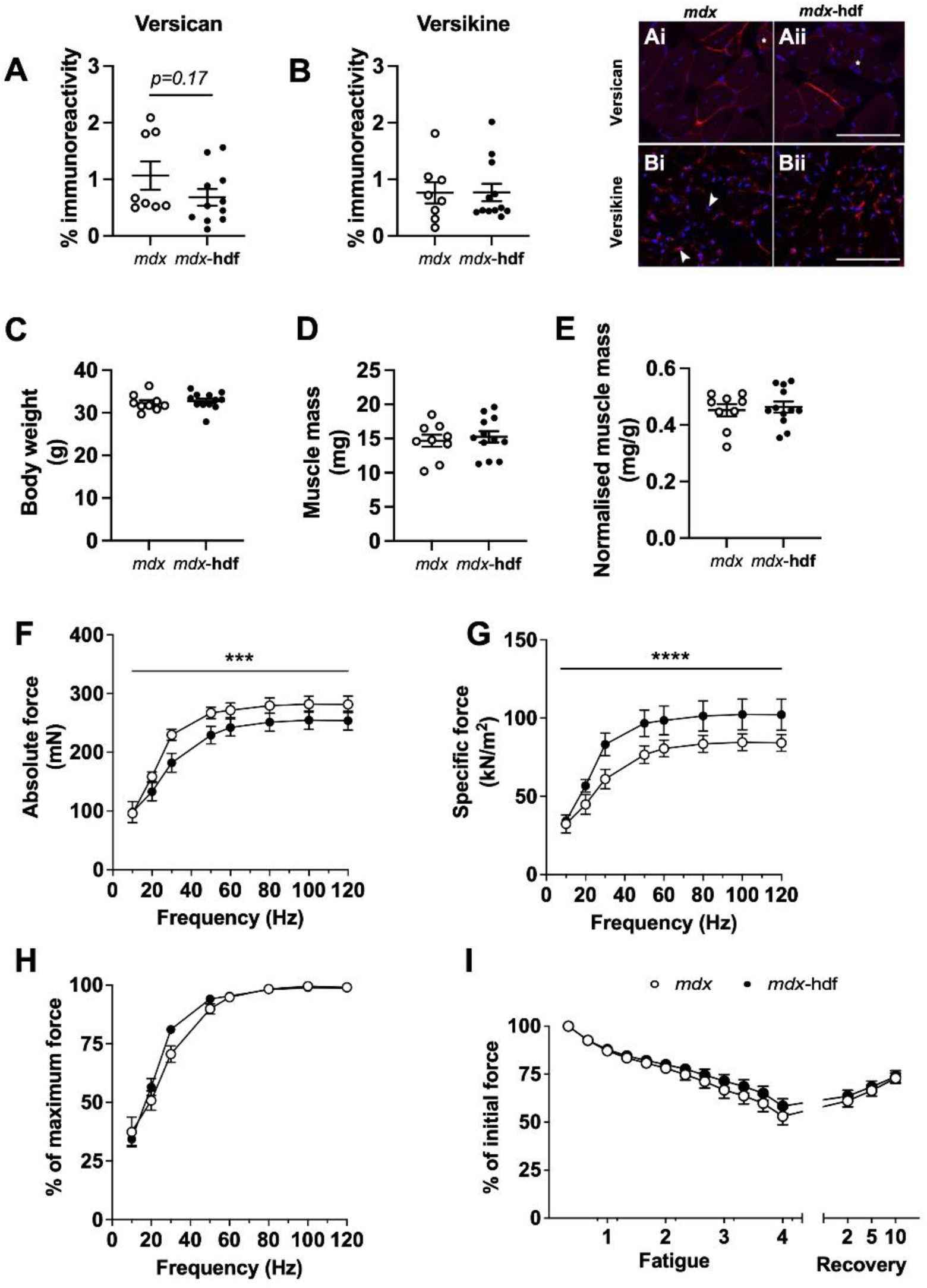
V0/V1 versican and versikine immunoreactivity and *ex vivo* strength and endurance of soleus muscles from adult *mdx* and *mdx*-hdf mice. Quantification of (A) versican and (B) versikine immunoreactivity. Representative images of versican or versikine staining (red) in soleus muscles from *mdx* (Ai, Bi) and *mdx*-hdf mice (Aii, Bii), nuclei are counterstained with DAPI (blue); scale bar = 100 μm. (C) Body mass and (D) the absolute and (E) normalized to body weight (mg of muscle mass per g of body weight) of soleus muscle from 20-week-old *mdx* and *mdx*-hdf mice. Force frequency curves expressed as (F) absolute force (P_o_), (G) specific force normalized to muscle cross-sectional area (sP_o_), and as (H) a percentage of maximal force output of soleus muscles. (I) Muscle fatigability and force recovery following 4 min of intermittent stimulation (60 Hz, every 5 s) expressed as a percentage of initial force output. ***P < 0.001; ****P < 0.0001; 2-way ANOVA – main effect genotype. *N* = 11-13 per group for immunohistochemistry and *N =* 9-12 per group for contractile function.

At 20 weeks of age, body weight and absolute or normalized to body weight soleus mass did not significantly differ between *mdx*-hdf mice and *mdx* littermates (Fig. 8C-E). In contrast to the increase in strength observed in soleus muscles from 6-week-old *mdx*-hdf mice (Fig. 5D and E), a downward shift in absolute and specific force produced across the force frequency curve was observed in soleus muscles from adult *mdx*-hdf mice (P = 0.001 and P < 0.0001, respectively; Fig. 8F and G). This suggests that these muscles were weaker when compared to *mdx* littermates. When force frequency curve was expressed as a percentage of maximal force output, versican haploinsufficiency did not result in a left or right axis shift (Fig. 8H). In addition, fatigability and force recovery did not significantly differ between soleus muscles from adult *mdx*-hdf mice and *mdx* littermate controls (Fig. 8I).

When compared to soleus muscles from 6-week-old *mdx*-hdf and *mdx* mice, by approximately 20 weeks of age both genotypes had undergone dramatic morphological changes indicative of more severe dystrophic pathology including the presence of vacuolated myofibers and ‘ghost fibers’ characterized by the complete loss of cytoplasmic and my ofibrillar protein (Fig. 9 Ai-Avi). Soleus muscles from *mdx*-hdf mice had a greater percentage of smaller sized fibers between 30-39.99 µm in diameter (*P* < 0.05) and fewer large fibers >70 µm in diameter (P < 0.05; Fig. 9A). Furthermore, soleus muscle cross-sections from *mdx*-hdf mice tended to have a greater percentage of mononuclear infiltrate and degeneration compared to *mdx* littermates (*P* = 0.09; Fig. 9B). These observations, together with the observed reduction in muscle force production, suggest that versican haploinsufficiency exacerbated soleus muscle pathology in adult *mdx* mice. The underlying mechanisms remain to be determined, as no differences were detected between genotypes in select regeneration and inflammation markers. Specifically, the percentage of centrally nucleated myofibers (Fig. 9C) and the number of desmin positive muscle progenitor cells (Fig. 9D) or CD68 positive monocytes and macrophages (Fig. 9E) per mm^2^ of muscle cross-section did not significantly differ between soleus muscles from *mdx*-hdf mice and *mdx* littermate controls.

**Figure 9.**
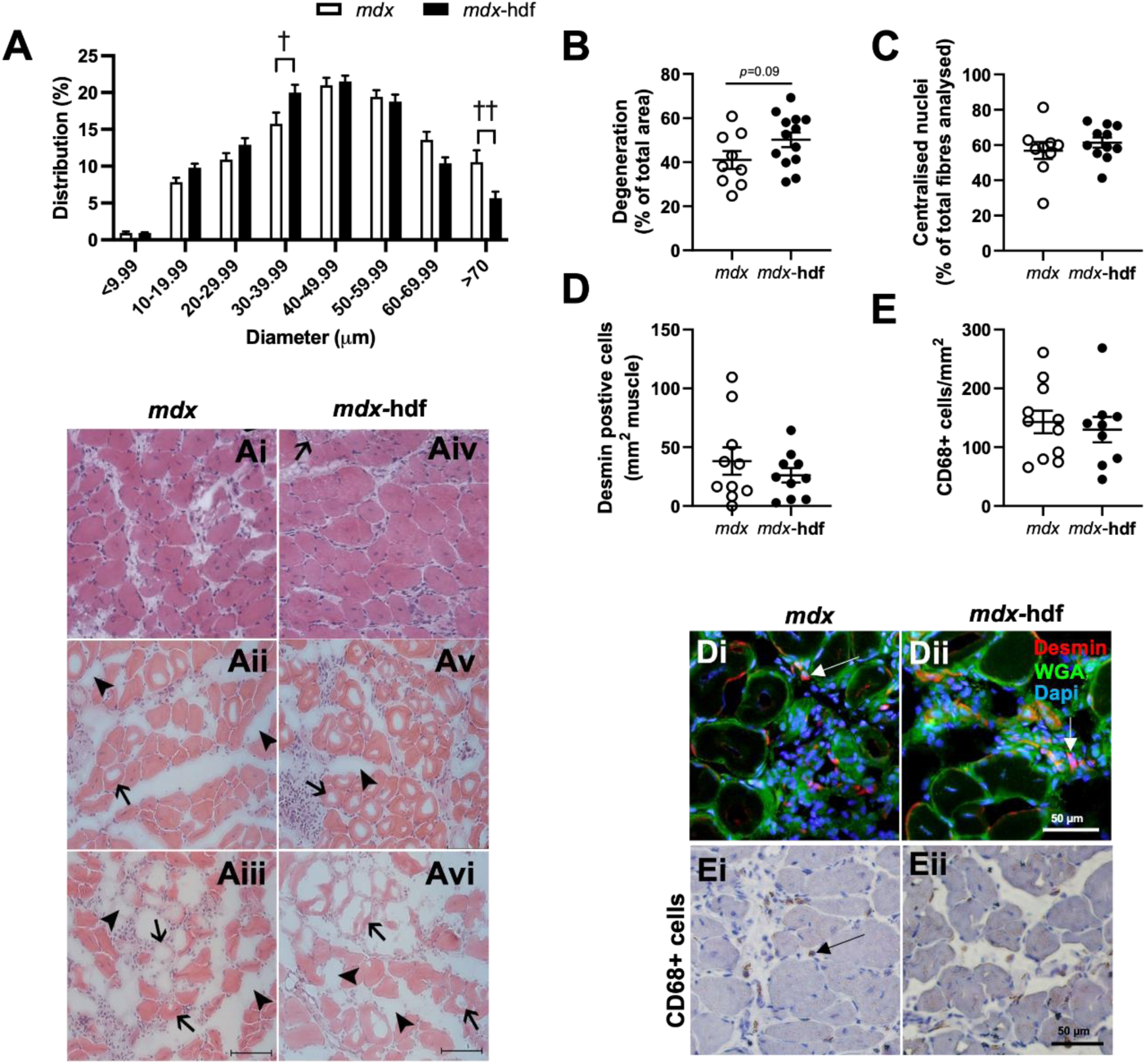
Muscle pathology and markers of inflammation and regenerative myogenesis in soleus muscles from adult *mdx-*hdf and *mdx* mice. (A) Myofiber size distribution. Representative images of soleus muscle cross-sections with a moderate dystrophic pathology with intact undamaged muscle fibers, centrally nucleated muscles fibers, and mononuclear infiltrates (Ai and Aiv); with a more severe pathology; including degenerating fibers with vacuoles (black arrows) and ghost fibers (empty spaces surrounding by basement membrane; black arrowheads), and mononuclear infiltrates (Aii and Av); and with the most severe pathology characterized by mononuclear infiltrates, extensive myofiber degeneration, and ghost fibers (Aiii and Avi). Scale bar = 100 µm. (B) The percentage of muscle cross-section comprised of mononuclear infiltrates and degenerating tissue. (C) The percentage of centrally nucleated fibers. (D) Quantification of the number of desmin positive muscle progenitor cells. (E) CD68 positive monocytes and macrophages per mm^2^ of muscle. Representative images of desmin positive cells (red; white arrow) from *mdx* (Di) and *mdx-*hdf (Dii) mice with extracellular matrix and sarcolemma stained with WGA (green) and nuclei counterstained with DAPI; scale bar = 50 μm. Representative images of CD68 positive cells stained brown with 3,3’-diaminobenzidine (DAB; black arrow) from *mdx* (Ei) and *mdx-*hdf (Eii) mice with nuclei counterstained with haematoxylin; scale bar = 50 μm. ^†^ P < 0.05; ^††^ P < 0.01; independent T-test. *N* = 11-13 per group for histology and immunohistochemistry and *N =* 9-12 per group for gene expression analysis.

## DISCUSSION

The transient upregulation of provisional matrix components, such as versican, is increasingly recognized as being important for muscle repair, whereas a persistent upregulation is associated with fibrosis and heightened inflammation (25, 56, 58). Versican expression is increased in diaphragm and TA muscles from *mdx* mice versus wild type mice and is suppressed by glucocorticoids during myogenic differentiation *in vitro* (32). Building on these observations, we report that in hindlimb muscles from juvenile *mdx* mice, greater V0/V1 versican immunoreactivity was observed in slow soleus compared to fast TA and EDL muscles. Also, the glucocorticoid – α- methylprednisolone – decreased *V0/V1 versican* mRNA transcripts in soleus, but not EDL, muscles. In dystrophic hindlimb muscles, an association between muscle fiber type and versican appears to exist. Fiber type dependent differences in proteoglycan content are not well characterized; although, there is more collagen slow versus fast muscles (59, 60). Therefore, studies investigating ECM components in dystrophic muscles need to consider the potentially confounding effects of muscle phenotype and age. We have previously shown that in fibrotic diaphragm muscles from adult (20- to 26-week-old) *mdx*-hdf mice, versican haploinsufficiency attenuated inflammation and improved contractile function, and this was associated with increased spontaneous physical activity (24). Unlike in the diaphragm, muscle fibrosis is minimal in hindlimb muscles from *mdx* mice (61). Here we report, that in juvenile and adult *mdx*-hdf mice, fiber type and age modulate the effects of versican haploinsufficiency on contractile function and importantly that in the absence of fibrosis, inflammation is not attenuated.

The hdf mouse has been used extensively to investigate the cellular mechanisms mediated by the genetic reduction of versican in various developmental processes and disease models. These include the transition of dermal fibroblasts to myofibroblasts (62), palatogenesis and cleft palate malformation (63), cardiac cushion remodeling during heart valve development (64), angiogenesis and tumor growth in cancer (65), and T cell immunity during influenza virus infection (66). In 6-week-old *mdx* mice, the effect of versican haploinsufficiency on versican mRNA transcript abundance and immunoreactivity varied with hindlimb muscle phenotype. A more robust decrease in versican expression was achieved in TA and soleus versus EDL muscles. The decrease in versican expression in soleus muscles from *mdx*-hdf mice appeared to be transient and may be associated with ECM remodeling during postnatal growth and/or the bout of spontaneous hindlimb muscle degeneration (37). At 20 weeks of age, soleus pathology was characterized by extensive myofiber degeneration with limited fibrosis, and versican expression did not significantly differ between *mdx*-hdf and *mdx* mice.

In soleus muscles from 6-week-old *mdx* mice, versican reduction was associated with increased absolute (P_o_) and specific (sP_o_) muscle force output and a modest reduction in fatiguability. This reduction in fatigability was also observed in EDL muscles. Notably, similar positive effects on *ex vivo* contractile function were observed in diaphragm muscles from adult *mdx*-hdf mice. However, unlike in the diaphragm, this improvement in soleus muscle contractile function with versican haploinsufficiency was not sustained into adulthood. Soleus muscles from 20-week-old *mdx*–hdf mice produced less force output compared to their *mdx* littermates, as indicated by a downward shift in the absolute (P_o_) and specific (sP_o_) force frequency curves at 20 weeks of age. We hypothesize that these functional differences between soleus and diaphragm muscles from adult *mdx*-hdf mice may be mediated by the low level of fibrosis in the soleus muscle and indicate that a threshold level of versican expression is needed for optimal muscle physiology.

The negative consequences of versican haploinsufficiency on soleus muscle contractile function in 20-week-old *mdx*-hdf mice were unexpected; especially, given previous observations of increased spontaneous physical activity (24). Versican haploinsufficiency also tended to exacerbate soleus muscle degeneration and had deleterious effects on myofiber size. Although, a profound deterioration of soleus muscle architecture was observed irrespective of genotype. This is corroborated by Pastoret *et al.* who described more degenerating and regenerating fibers in soleus than EDL muscles from *mdx* mice at 13 and 26 weeks of age (67). As well as by Chamberlain *et al.* who reported that the degree of dystrophic pathology is not a simple reflection of fiber type and that soleus muscles from 26-month-old *mdx* mice presented with an extensive dystrophic pathology intermediate in severity between the diaphragm and hindlimb muscles with a faster phenotype (*e.g.*, quadriceps and TA) (68). Whilst fast twitch dystrophic muscle fibers are more susceptible to eccentric contraction induced damage (69) , the soleus undergoes far greater (∼3-fold) muscle excursion — the change in muscle length required to produce the full range of joint motion — compared to the EDL or TA and this may lead to a greater pathology (70). Altogether, our *ex vivo* contractile function experiments highlight the importance of muscle phenotype and age when investigating the role of versican in dystrophic *mdx* mice.

Biological context is also emerging to be important determinant of the immunomodulatory properties of versican (71). The hypothesis that a threshold level of versican expression is required for optimal muscle function may also apply to its pro-inflammatory properties in dystrophic muscles. In fibrotic diaphragm muscles from adult *mdx*-hdf mice, versican reduction reduced the infiltration of CD68 positive monocytes and macrophages by ∼50%, whereas in soleus muscles the number of CD68 positive inflammatory cells did not significantly differ between juvenile or adult *mdx*-hdf mice and *mdx* littermates. A limitation of using 6-week-old *mdx*-hdf mice, is that the effect of versican haploinsufficiency on muscle inflammation during the peak hindlimb degeneration phase is not known. Peak degeneration occurs in *mdx* hindlimb muscles at 3-4 weeks of age and is characterized by extensive provisional ECM synthesis – including fibronectin and tenascin-C, known versican binding partners (15, 72, 73).

Versican may have a role in regulating the ECM microenvironment of the satellite cell niche to favor activation and proliferation. Aligned with a high level of muscle regeneration, versican gene expression is highly upregulated in isolated satellite cells from 6-week-old *mdx* mice compared to age matched wild type mice (74). We did not detect an effect of versican haploinsufficiency on the markers of muscle regeneration assessed. *MyoD* and *Myogenin* mRNA transcripts, desmin positive muscle progenitor cells or centrally nucleated fibers did not significantly differ in soleus muscles from 6-week-old or 20-week-old *mdx*-hdf and *mdx* mice. However, a limitation of our study is that effect of versican haploinsufficiency on satellite cell activation and the consequences on myoblast proliferation and differentiation were not comprehensively investigated. The role of ECM proteins in regulating regenerative myogenesis, including muscle progenitor cell behavior, is increasingly recognized, as is the concept that dysregulation of ECM components will compromise regenerative myogenesis (75). There are interesting parallels to be drawn between versican and fibronectin, a provisional matrix protein also upregulated in dystrophic muscles and associated with fibrosis (76, 77). Like versican, fibronectin has a dual role in inflammation (72) and muscle regeneration (25). Within hours of activation, muscle satellite cells from healthy muscles begin to secrete significant amounts of fibronectin (78). This autologous expression of fibronectin is required for satellite cell proliferation, as knockdown of fibronectin in a cardiotoxin injury model of muscle regeneration led to a dramatic decrease in satellite cell number (78).

We observed a reduction in fatigability of the EDL and soleus muscles of 6-week-old *mdx*-hdf mice; although, the effect was greater in the EDL. Similar effects on muscle fatigability were also observed in diaphragm muscles from adult (20- to 26-week-old) *mdx*-hdf mice and the genetic reduction of versican also modulated whole body metabolism leading to a reduction in glucose oxidation (24). Using immunohistochemical and histological staining for myosin heavy chain isoforms and succinate dehydrogenase (SDH) activity, we did not detect a significant effect of versican haploinsufficiency on muscle fiber type which could account for the improvement in hindlimb muscle endurance *ex vivo*. Further studies are needed to better understand how versican might modulate endurance and oxidative metabolism in dystrophic muscles. Impaired mitochondrial function is closely associated with increased oxidant production which are thought to drive DMD pathology secondary to the loss of dystrophin (79–82). ECM-mitochondria crosstalk is increasingly recognized as important for optimal organelle function (83). Indeed, there is evidence that skeletal muscle mitochondrial function is regulated by signals from the ECM. For example, TGFẞ is a positive regulator of Mss51 and increased Mss51 expression compromises mitochondrial function via complex IV (84). In *mdx* mice, the genetic deletion of Mss51 increased the mitochondrial respiratory capacity of isolated extensor digitorum brevis myofibers and reduced diaphragm histopathology – including markers of fibrosis (85). Further highlighting the potential significance of versican to skeletal muscle metabolic health, single-nucleotide polymorphisms in versican (*VCAN*; rs2287926) have been found to be to be associated with lean mass and lean appendicular mass, as well as metabolic protection (86, 87). More broadly, there is increasing interest in understanding genetic modifiers of DMD pathology (88) – and in that regard *VCAN* may be a significant, novel modifier.

In conclusion, we report that versican haploinsufficiency has phenotype and age specific effects on versican expression and muscle contractile function in hindlimb muscles of juvenile and adult *mdx* mice which are independent of changes in inflammation. This contrasts with our previously published findings in diaphragm muscles of *mdx*-hdf mice, where we showed an association between versican reduction and decreased macrophage and monocyte infiltration (24). We suspect this discrepancy is likely mediated by the low level of fibrosis in hindlimb versus diaphragm muscles from *mdx* mice (61). These data in *mdx*-hdf mice and that *V0/V1 Versican* mRNA transcripts were decreased by glucocorticoids in soleus, but not EDL, muscles of juvenile *mdx* mice, highlight the importance of muscle selectivity when investigating mechanisms mediating inflammation and fibrosis in dystrophy. There is an unmet need for effective anti-inflammatory and anti-fibrotic therapies for DMD. This is predicated upon an improved understanding of the interdependence between the ECM and the immune system in dystrophic muscles. We recommend future studies carefully consider the differences in the muscle pathology, fiber type and ECM composition as responses to an intervention may differ.

## ACKNOWLEDGMENTS

The authors would like to thank A/Prof Emma Rybalka (Victoria University) for the thoughtful reading of the manuscript and her insightful feedback on the discussion.

## AUTHOR CONTRIBUTIONS

Correspondence: N Stupka

## AUTHOR NOTES

Conceived and designed research: NS, DMC.

Performed experiments, analyzed data, interpreted results of experiments —NMR, DD, NS, AA, DMC, RB, RM, AH.

Prepared figures, drafted manuscript – DD, NS. Edited and revised manuscript – DD, NS, AA, NMR.

Approved final version of manuscript —NMR, DD, NS, AA, DMC, RB, RM, AH.

